# Dynamic *S*-acylation of GSDMA regulates pyroptosis

**DOI:** 10.1101/2025.10.16.682910

**Authors:** Zhipeng Tao, Ritesh P. Thakare, Melyssa Cheung, Jianan Zhang, Xin Chen, Xiaodi Hu, Brian G. Pierce, Xu Wu, Junhao Mao

**Affiliations:** Cutaneous Biology Research Center, Massachusetts General Hospital, Harvard Medical School, Charlestown, MA 02129, USA; Department of Molecular, Cell and Cancer Biology, University of Massachusetts Chan Medical School, Worcester, MA 01605, USA; Department of Chemistry and Biochemistry, University of Maryland, College Park, MD 20742, USA; Department of Cell Biology and Molecular Genetics, University of Maryland, College Park, MD 20742, USA; University of Maryland Institute for Bioscience and Biotechnology Research, Rockville, MD 20878, USA; Department of Nutrition and Food Sciences, Texas Woman’s University, Denton, TX 76204, USA

## Abstract

GSDMA, the primary member of the gasdermin family found in the skin, is critical for pathogen-induced pyroptosis during infection. Recent studies revealed that another gasdermin, GSDMD, undergoes palmitoylation during pyroptosis. However, whether and how the other gasdermin members undergo lipid modification remains poorly understood. Here, we demonstrate that GSDMA is *S*-acylated at the conserved cysteine residues in its N-terminal domain. We show that *S*-acylation of GSDMA promotes pyroptosis by facilitating its membrane anchoring and protein oligomerization, a mechanism distinct from the palmitoylation of GSDMD at the N-terminal C191 residue. Additionally, we present evidence that recombinant proteins of GSDMA and GSDMD can undergo *S*- acylation *in vitro* via direct interaction with palmitoyl-CoA, suggesting they potentially possess auto-acylation capacity. Furthermore, we identify ABHD17A as one of the deacylating enzymes that regulate the dynamic fatty acylation cycle of GSDMA. Overall, our studies reveal new molecular mechanisms underlying GSDMA function through S-acylation and underscore its important role in regulating pyroptosis mediated by GSDMA.

**Highlights:**

- GSDMA is *S*-acylated at N-terminal conserved cysteine residues.
- GSDMA *S*-acylation facilitates membrane anchoring and protein oligomerization.
- GSDMA and GSDMD recombinant proteins can undergo *S*-acylation in vitro
- The ABHD17A enzyme is involved in GSDMA deacylation.

## Introduction

Gasdermin A (GSDMA) is a member of the gasdermin family proteins, including GSDMB, GSDMC, GSDMD, and GSDME, which orchestrates pyroptosis, a type of inflammatory programmed cell death^1–6^ . GSDMA is predominantly expressed in keratinocytes and activated through cleavage by group A Streptococcus (GAS) cysteine protease, specifically by the virulence factor SpeB^7, 8^. This cleavage releases its N-terminal domain, which then forms pores in the cell membrane and initiates lytic cell death during skin infection^7–10^ . Thus, it is crucial to understand the mechanisms regulating GSDMA- mediated pyroptosis.

Palmitoylation is a form of post-translational fatty acylation involving the reversible attachment of a 16-carbon palmitoyl group to specific cysteine residues of proteins ^11–13^. Recent studies revealed that GSDMD undergoes S-palmitoylation at Cys191, likely catalyzed by palmitoyl acyltransferases, which promotes the association of GSDMD with the cell membrane, thereby facilitating pyroptosis^14–20^. Several small molecule inhibitors targeting GSDMD palmitoylation have also been identified, including Disulfiram, Dimethyl fumarate (DMF), and NU6300^21–23^. These compounds are thought to specifically covalently modify Cys191 of human GSDMD, and impair its palmitoylation, ultimately leading to pyroptosis inhibition^21–24^. These studies highlight the potential of targeting GSDMD palmitoylation as a therapeutic strategy for inflammatory diseases. However, Cys191 is not conserved in the other members of the gasdermin family. Thus, whether and how the other gasdermin proteins undergo lipid modification during pyroptosis remains largely unknown.

In this study, we demonstrated that GSDMA is *S*-acylated at the conserved cysteine residues in its N-terminal domain, distinct from the reported Cys191 palmitoylation in GSDMD. We showed that the recombinant GSMDA and GSDMD proteins could undergo acylation *in vitro* independent of palmitoyl transferases by direct interaction with palmitoyl-CoA. We also presented evidence that the dynamic GSDMA acylation is partly mediated by the deacylating enzyme ABHD17A, providing a potential new regulatory mechanism of GSDMA during pyroptosis.

## Results

### Dynamics of GSDMA3 fatty acylation

To examine whether GSDMA undergoes fatty acylation, we utilized a series of fatty acid chemical probes with different chain lengths, including 14-carbon myristoylation probe (Alk-C14), 16-carbon palmitoylation probe (Alk-C16), 18-carbon stearoylation probe (Alk- C18), 20-carbon arachidic acid (Alk-C20) (Figure 1A). Flag-tagged mouse GSDMA3 was transfected into the HEK293A cells, labeled cells with 50 µM of probes, followed by Click reaction with biotin-azide. The streptavidin blot results suggested that Alk-C16 and Alk- C18 probes displayed higher labeling efficiency, whereas Alk-C14 and Alk-C20 probes showed lower efficiency (Figure 1B). In contrast, we showed that the various fatty acid chemical probes could not label the control protein YAP in transfected HEK293 cells (Figure S1A). Together, these data demonstrated that GSDMA3 could be fatty acylated and GSDMA3 fatty acylation prefers 16-carbon palmitic acid and 18-carbon stearic acid. To test whether endogenous GSDMA protein can be acylated, COLO205 cells were metabolically labeled with 100 µM Alk-C16, followed by a Click reaction with biotin-azide. The fatty acylated proteome was then enriched by streptavidin beads pull-down. Western blotting for GSDMA in the pull-down samples indicated that endogenous GSDMA was indeed acylated in cells (Figure 1C).

**Figure 1.**
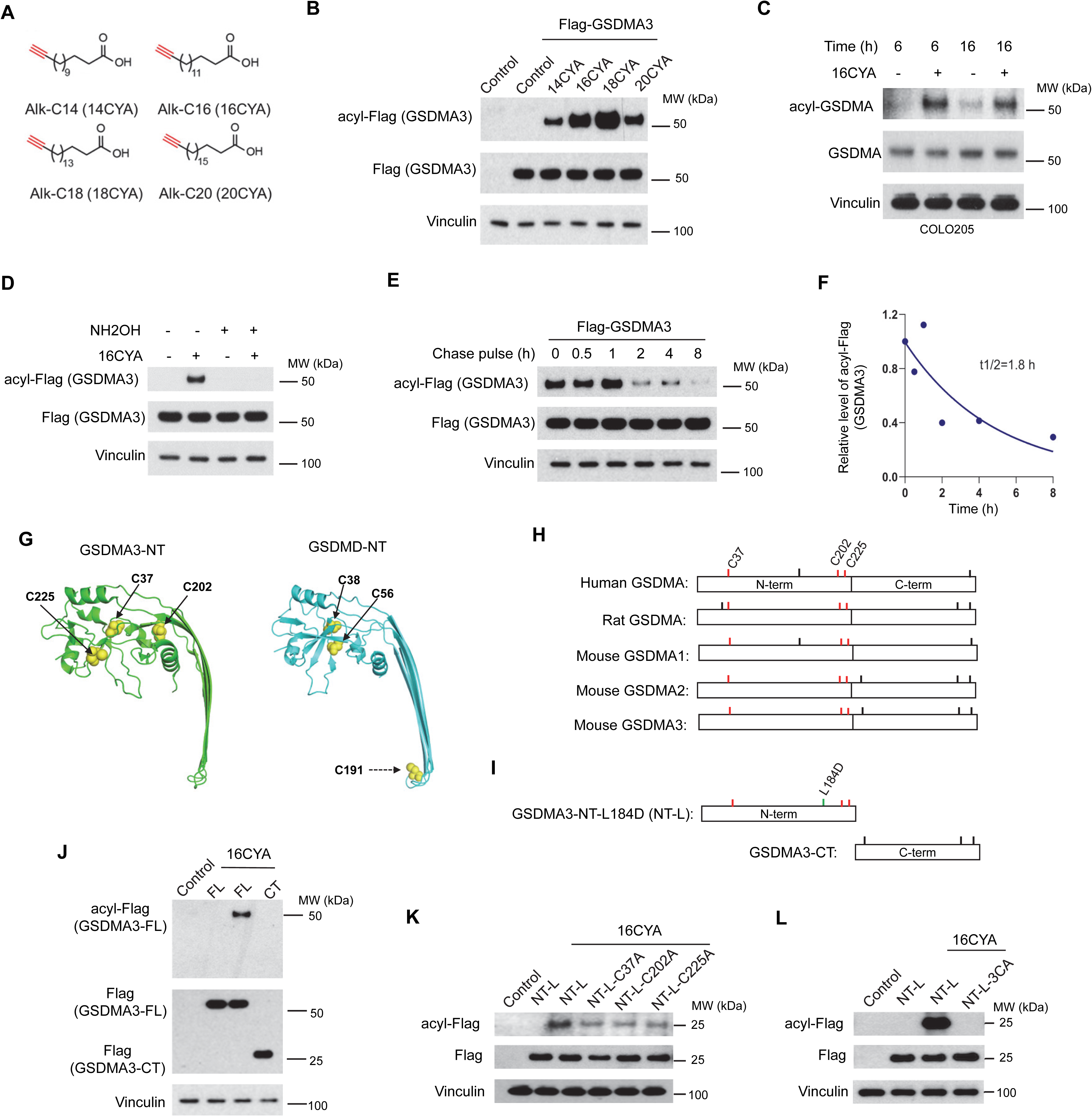
Dynamic acylation of GSDMA3 at conserved cysteines in the N-terminal domain. (A) Different chain length fatty acid probes; (B) *S*-acylation of GSDMA3 in transfected HEK293 cells measured by different chain length fatty acid probes; Flag (GSDMA3) represents the input control; acyl-Flag (GSDMA3) represents the amount of GSDMA3 been lapidated. (C) Fatty acylation of endogenous GSDMA in colorectal cancer COLO205 cells; (D) Acylation of GSDMA3 is hydrolyzed by 0.5% hydroxylamine (NH_2_OH) (v/v); (E and F) The dynamic and half-life of GSDMA3 *S*-acylation. The half-life is 1.8 hours (F); (G) The structures of GSDMA3 and GSDMD N-terminal domains (GSDMA3- NT and GSDMD-NT), from PDB codes 6CB8 and 6VFE, respectively (with GSDMA3-NT missing residues modeled as noted in Methods), with their cysteine residues labeled; (H) The schematic representations of GSDMA proteins. The conserved N-terminal cysteine residues C37, C202 and C225 are labeled in red; (I) The schematic representations of GSDMA3 N-terminal domain with L184D mutation (NT-L) and C-terminal domain (CT). (J) Acylation of full-length (FL) and C-terminal (CT) GSDMA3 in transfected HEK293 cells; (K and L) Acylation of GSDMA3-NT-L carrying C37A, C205A or C225A mutation (K), or all three cysteines mutated to alanine (NT-L-3CA) (L).

In addition, we found that the treatment of Click reaction products with 2.5% (vol/vol) hydroxylamine (NH_2_OH) abolished the acylation signal of GSDMA3 in transfected HEK293 cells, confirming that GSDMA3 is S-acylated through thioester bonds with cysteine residues (Figure 1D). Next, we tested whether GSDMA3 fatty acylation is dynamically regulated through acylation and deacylation cycles. We detected GSDMA3 acylation after 1-hour labeling with Alk-C16 and the signal increased further up to 12-hour of labeling (Figure S1B). To determine the rates of fatty acylation cycling of GSDMA3, pulse-chase experiments were performed in transfected HEK293 cells, and the results showed that the turnover rate of GSDMA3 acylation is approximately 1.8 h, while the protein remained stable during the assay period (Figure 1E and 1F). These results indicate that *S*-acylation of GSDMA3 is a dynamic process.

### GSDMA3 is S-acylated at conserved cysteine residues in the N-terminal domain

Next, we set out to determine the acylation site(s) of GSDMA. Several recent reports showed that GSDMD is palmitoylated at Cysteine 191 (C191) in the N-terminal domain^14–17, 23^ . However, C191 is not conserved in the N-terminal domains of other gasdermin proteins, including GSDMA (Figure 1G, 1H, and 1I), suggesting that the other Cysteine residues may be responsible for palmitoylation.

We first showed that in comparison to the full-length protein, the C-terminal domain of GSDMA3 did not have a detected acylation signal when expressed in HEK293 cells, suggesting that, similar to GSDMD, the N-terminal domain of GSDMA undergoes lipid modification (Figure 1J). We then aligned the sequences of GSDMA proteins across different species, including human and rat GSDMA and mouse GSDMA1/2/3, and found three conserved cysteine residues (C37, C202 and C225) in the N-terminal domain (Figure 1H). To identify the acylated cysteine residues, we constructed a series of GSDMA3 N-terminal mutants in which the three cysteine residues were mutated to alanine, and they also carried a leucine 184 to aspartic acid (L184D) mutation that has been shown to suppress partially pyroptosis allowing sufficient expression of the N-terminal domain of GSDMA3 in cells for out click-chemistry and pull-down assays^25^ (Figure 1I). Our results showed that the three cysteines can undergo acylation, as the individual C37A, C205A, or C225A mutants reduced the signal levels (Figure 1K); however, the mutation of all three cysteine residues could completely abolish GSDMA acylation in the N-terminal domain (Figure 1L).

### GSDMA and GSDMD recombinant proteins undergo *S*-acylation *in vitro*

A recent report showed that bacterial gasdermin can undergo auto-palmitoylation^26^. Despite the sequence and structure differences, this raises the intriguing question of whether mammalian gasdermin proteins can also undergo acylation independent of palmitoyl transferases by direct interaction with palmitoyl-CoA.

Several recent studies revealed that the palmitoylation of GSDMD at its N-terminal C191 residue is mediated by ZDHHC5, 7, or 9 palmityl acyltransferases. Interestingly, we found that ectopic expression of a collection of ZDHHC enzymes, including ZDHHC5, 7, or 9, did not significantly increase the acylation levels of GSDMA3 in cells (Figure S1C, D). To further examine ZDHHC5, 7, 9, the three ZDHHC enzymes involved in GSDMD palmitoylation, we silence their expression in cells using shRNA (Figure S1E-G). We found that the knockdown of these ZDHHC enzymes did not affect GSDMA3 acylation (Figure S1H). These data suggested that the ZDHHC enzymes involved in GSDMD palmitoylation may not play a significant role in GSDMA acylation.

To further test whether GSDMA can be acylated independent of palmitoyl transferases by direct interaction with palmitoyl-CoA, we used the recombinant human GSDMA protein in biochemical assays *in vitro*. GSDMA protein was incubated with physiologically relevant concentrations (1uM, 2.5 μM, 5 μM, or 10 μM) of a clickable analog of palmitoyl-CoA for 30 min at neutral pH, followed by Click reaction with biotin-azide. The streptavidin blot showed that GSDMA was strongly acylated *in vitro* in a palmitoyl-CoA concentration-dependent manner (Figure 2C). Next, we fixed the concentration of palmitoyl-CoA at 5 μM and examined GSDMA acylation at different time points. Our results suggested that the acylation level of GSDMA increases in a time-dependent manner (Figure 2D). Furthermore, we characterized the GSDMA concentration and temperature dependent reactions with palmitoyl-CoA. We found that GSDMA reacts with palmitoyl-CoA in a temperature dependent manner (Figure S2A) and that the reaction between palmitoyl-CoA and GSDMA also depends on the concentration of GSDMA, reaching saturation at a concentration of 500 nM (Figure S2B). In addition, we showed that treating the Click reaction products with 2.5% (vol/vol) NH_2_OH effectively abolished GSDMA acylation, suggesting the *in vitro* S-fatty acylation of recombinant GSDMA is sensitive to NH_2_OH (Figure 2B). To calculate the kinetic of GSDMA’s enzymatic reaction, we determined the initial velocity (V0) of GSDMA *in vitro* acylation reactions under different concentrations of palmitoyl-CoA when the reaction time was fixed at 15 minutes. The apparent Km of palmitoyl-CoA on GSDMA3 palmitoylation is around 3.8 μM (Figure 2E and 2F), which is comparable to the Km of ZDHHC-PATs ^27, 28^. The Kcat value is 4.2 × 10^7^ /min, while Kcat/Km is 1.1 × 10^13^ L/min*mol. Together, these data demonstrated that the GSDMA recombinant protein can undergo *S*-acylation *in vitro*.

**Figure 2.**
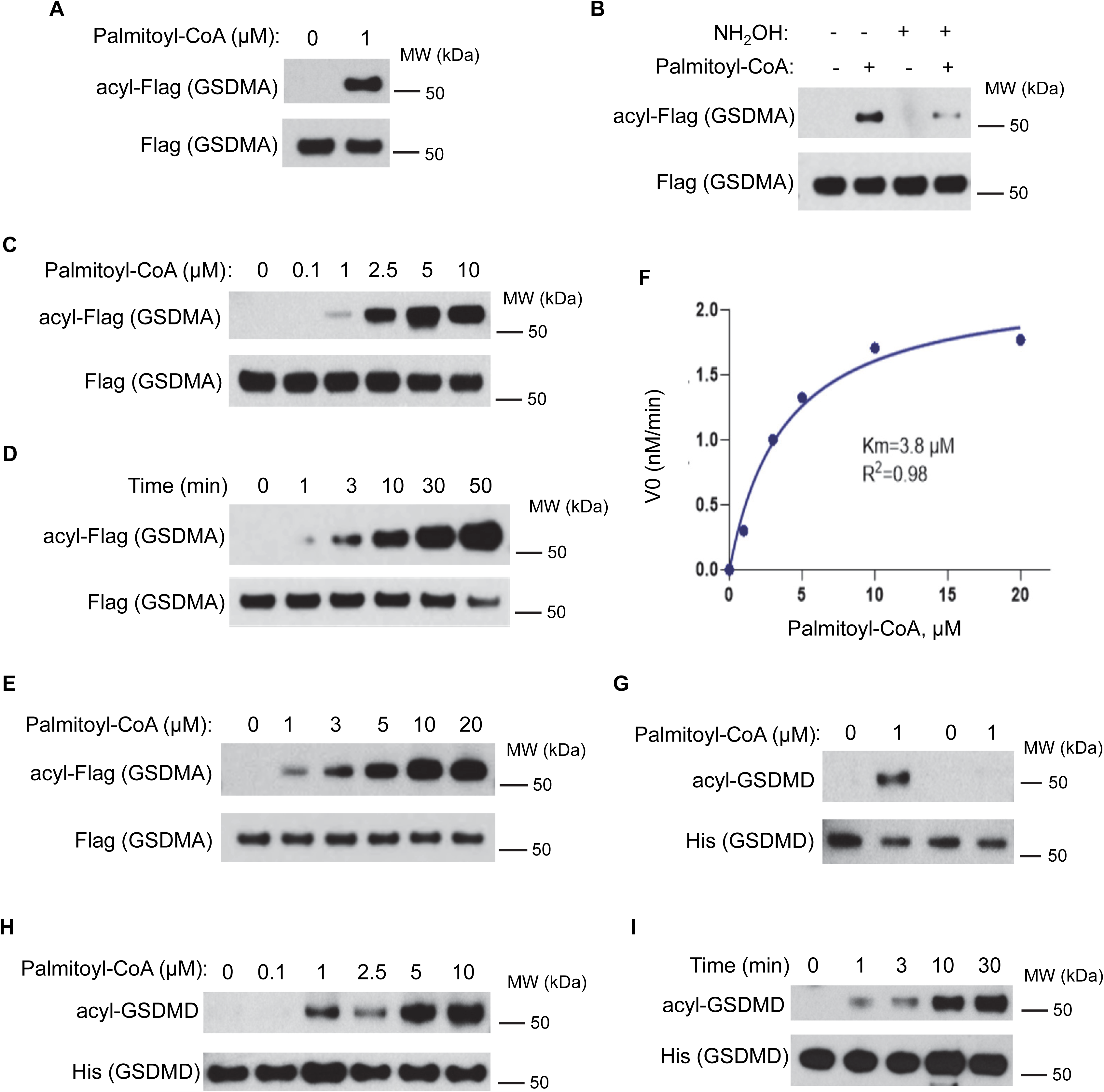
Acylation of gasdermin proteins *in vitro*. (A) Recombinant GSDMA protein reacts with alkyne-palmitoyl-CoA (1 µM) *in vitro* independent of palmitoyl acyltransferases; (B) Acylation of recombinant GSDMA by alkyne-palmitoyl-CoA (1 µM) was significantly reduced by NH_2_OH; (C) Acylation of recombinant GSDMA with different concentrations of alkyne-palmitoyl-CoA (0, 0.1, 1, 2.5, 5, and 10 µM); (D) Acylation of recombinant GSDMA with 1 µM alkyne-palmitoyl-CoA for different durations (0, 1, 3, 10, 30, and 50 minutes); (E) The reaction of recombinant GSDMA with different concentrations of alkyne-palmitoyl-CoA (0, 1, 3, 5, 10, and 20 µM) for 15 minutes to determine the *Km* value; (E) The plot of 3(E) and the calculation of the *Km* value of the reaction, Km= 3.8 µM and R^2^=0.98; (G) Acylation of recombinant GSDMD with alkyne-palmitoyl-CoA (1 µM) with or without NH_2_OH treatment; (H) Acylation of recombinant GSDMD with different concentrations of alkyne-palmitoyl-CoA (0, 0.1, 1, 2.5, 5, and 10 µM); (I) Acylation of recombinant GSDMD with 1 µM alkyne-palmitoyl-CoA for different durations (0, 1, 3, 10, and 30 minutes)

The ability of GSDMA to undergo acylation by direct interaction with palmitoyl-CoA prompted us to test the other members of the gasdermin family, including GSDMD. Interestingly, we found that the recombinant GSDMD protein also reacted with alkyne- palmitoyl-CoA *in vitro*, and its palmitoylation signal was also sensitive to NH_2_OH treatment (Figure 2G). Furthermore, we demonstrated that GSDMD could be acylated in a time-dependent and dose-dependent manner with different concentrations of alkyne-palmitoyl-CoA *in vitro* (Figure 2H and 2I). Taken together, these results suggested that the gasdermin family members can bind to palmitoyl-CoA and undergo *S*-acylation *in vitro*.

### Acylation of GSDMA3 is critical for membrane anchoring during pyroptosis

To gain insights into the structural basis of the potential function of GSDMA acylation, we generated a model of palmitoylated GSDMA3 N-terminal domain in a lipid membrane, using structural modeling to add palmitoylation at three cysteines and membrane to a previously determined structure of GSDMA3 protein pore1 (detailed in Methods)^29^ . The model (Figure 3A, 3B and 3C) shows that the C225 palmitate is positioned to interact with the GSDMA3 hydrophobic anchor and membrane lipids (Figure 3C). In contrast, the C37 and C202 palmitates are predicted to be less embedded in the membrane (Figure 3C). They are more heterogeneous in their positioning and orientations among the modeled pore subunits, possibly indicating less energetic favorability or a need for additional modeling to confirm their native conformations. Interestingly, we also found that all three palmitates are proximal to adjacent pore subunits (Figure 3D). Thus, our structural modeling suggested that *S*-acylation of GSDMA3 at C37, C202, and C225 in the N-terminal domain might mediate its membrane anchoring, and its potential involvement in the inter-unit interactions in the pore assembly.

**Figure 3.**
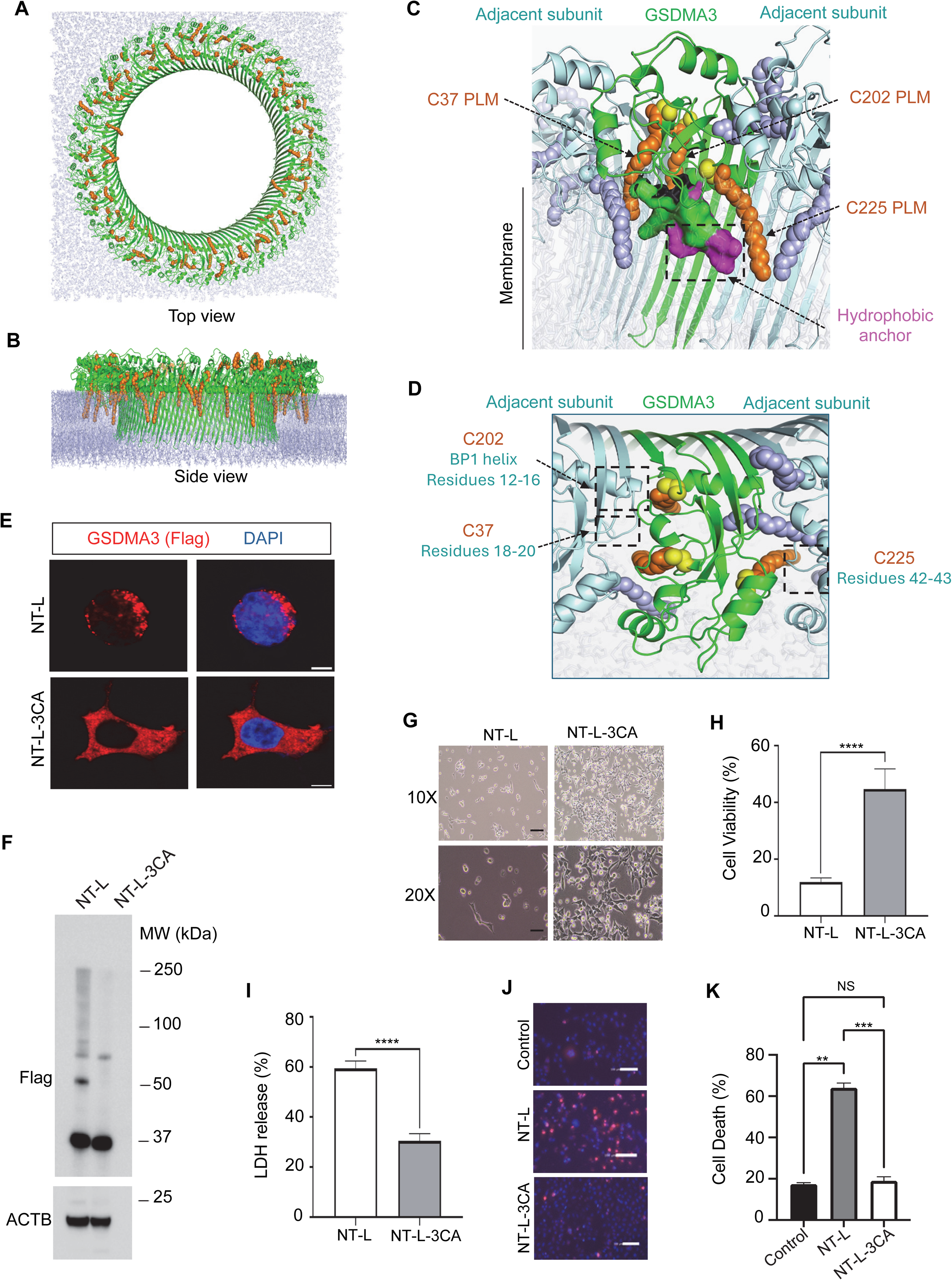
GSDMA3 acylation facilitates membrane anchoring and promotes pyroptosis. (A) Top view and (B) side view of membrane-bound GSDMA3 pore (green) with modeled palmitoylation (orange), and lipids shown as light blue sticks; (C) Side view of GSDMA3 protein (green) in pore with modeled palmitates (orange) and hydrophobic anchor (magenta) labeled, with adjacent subunits colored light blue, and membrane lipids shown as gray sticks; (D) Top view of GSDMA3 protein and adjacent subunits from (C), with proximal adjacent subunit residues for each modeled PLM noted; (E) Immunofluorescent staining of GSDMA3-NT-L with or without mutation of cysteine residues (NT-L-3CA) in transfected HEK293 cells. GSDMA3-NT-L and GSDMA3-NT-L- 3CA proteins carrying a C-terminal Flag tag were detected by an anti-Flag antibody; (F) Oligomerization of GSDMA3-NT-L and GSDMA3-NT-L-3CA in transfected HEK293 cells was measured by the semi-Native SDS-PAGE and detected by an anti-Flag antibody; (G) The representative images of HEK293 cells expressing GSDMA3 NT-L or NT-L-3CA; (H) Cell viability of HEK293 cells expressing GSDMA3 NT-L or NT-L-3CA; (I) LDH release from HEK293 cells expressing NT-L or NT-L-3CA; (J) Fluorescent images that visualize cell death in HaCat cells expressing NT-L or NT-L-3CA. Propidium iodide (PI) staining (red) highlights dead cells, while Hoechst 33342 staining (blue) marks cell nuclei. (K) Cell death in HaCat cells expressing NT-L or NT-L-3CA. n=3; **, p < 0.01; ***, p < 0.001; ****, p < 0.0001; NS, no significance.

To test these models, we first examined whether GSDMA acylation at its N-terminus is required for membrane localization. HEK293 cells were transiently transfected with GSDMA N-terminal domain carrying a leucine 184 to aspartic acid (L184D) mutation (NT- L) with or without the three cysteine residues were mutated to alanines (NT-L-3CA). Their localization was determined by immunofluorescence imaging (Figure 3E). Consistent with previous studies, GSDMA3-NT-L was predominantly membrane localized ^29^, whereas GSDMA3-NT-L-3CA was cytosolic (Figure 3E). Next, we investigated the effect of acylation on GSDMA-NT oligomerization. Western blotting revealed that GSDMA3-NT-L, but not the 3CA mutant, formed oligomers (Figure 3F). To determine whether GSDMA3 acylation modulates pyroptosis, we performed the LDH cytotoxicity assay and measured cell viability in HEK293 cells transfected with GSDMA3-NT-L constructs with or without cysteine to alanine mutations. We found that the mutation of all three cysteine residues significantly improved cell viability and decreased LDH release (Figure 3G, 3H, and 3I). To further test the effect of GSDMA3 acylation in a more relevant cell context, we utilized the HaCat cells, the immortalized keratinocyte cell line derived from adult human skin. Our data indicated that GSDMA3-NT-L, but not GSDMA3-NT-L-3CA, significantly induced cell death in HaCat keratinocytes (Figure 3J and 3K). Together, these findings corroborate the functional importance of *S*-acylation in GSDMA3 localization, pore formation, and its associated pyroptosis in cells.

### ABHD17A regulates GSDMA3 de-acylation and pyroptosis

Our data indicated that the fatty acylation of GSDMA3 is dynamically regulated within cells, with a half-life of approximately 1.8 hours (Figure 1E and 1F). The dynamics of protein acylation is controlled by deacylating enzymes, such as APTs and ABHDs^30, 31^. To further investigate GSDMA3 de-acylation, we screened various potential deacylating enzymes, including ABHD1 through ABHD17C, LYPLA1 (APT1), LYPLA2 (APT2), LYPLAL1, PPT1, and PPT2 (Figure 4A, S1I). Although variability in GSDMA3 acylation was observed with co-expression of ABHD family members, our data indicate that multiple enzymes could potentially de-acylate GSDMA3 when ectopically expressed in HEK293 cells (Figure 4A, S1I). We further analyzed ABHD17A, LYPLA1, and LYPLAL1 due to their robust de-acylation effect on GSDMA3 (Figure 4A, 4B). Considering the expression levels of the three enzymes in transfected cells, ABHD17A exhibited the strongest activity against GSDMA3 (Figure 4B). In addition, ABHD17A is highly expressed in skin keratinocytes (Protein Atlas), where GSDMA plays a critical role in mediating pyroptosis^8,9^. Thus, we concentrated our analysis on ABHD17A. First, we assessed the effect of ABHD17A on pyroptosis of HEK293 cells induced by GSDMA3 NT-L. Our results indicated that ABHD17A over-expression increased cell viability and reduced LDH release compared to cells expressing GSDMA3 NT-L alone (Figure 4C, 4D, and 4E). However, ABHD17A overexpression appeared to have a lesser effect than GSDMA3 NT- L-3CA (Figure 3), suggesting that other enzymes, such as LYPLA1 and LYPLAL1, may also play a role in regulating GSDMA3 deacylation. Next, we examined the role of endogenous ABHD17A on GSDMA3 acylation and pyroptosis, by using siRNA to knockdown ABHD17A in HEK293 cells. We found that ABHD17A knockdown increased GSDMA3 acylation (Figure 4F) and exacerbated GSDMA3 NT-L-induced pyroptosis, measured by LDH release and cell viability (Figure 4G 4H and 4I). Taken together, our data suggest that GSDMA3 can be de-acylated by several enzymes, including ABHD17A, which together likely mediate the dynamic regulation of the acylation-deacylation cycle of GSDMA3 in cells.

**Figure 4.**
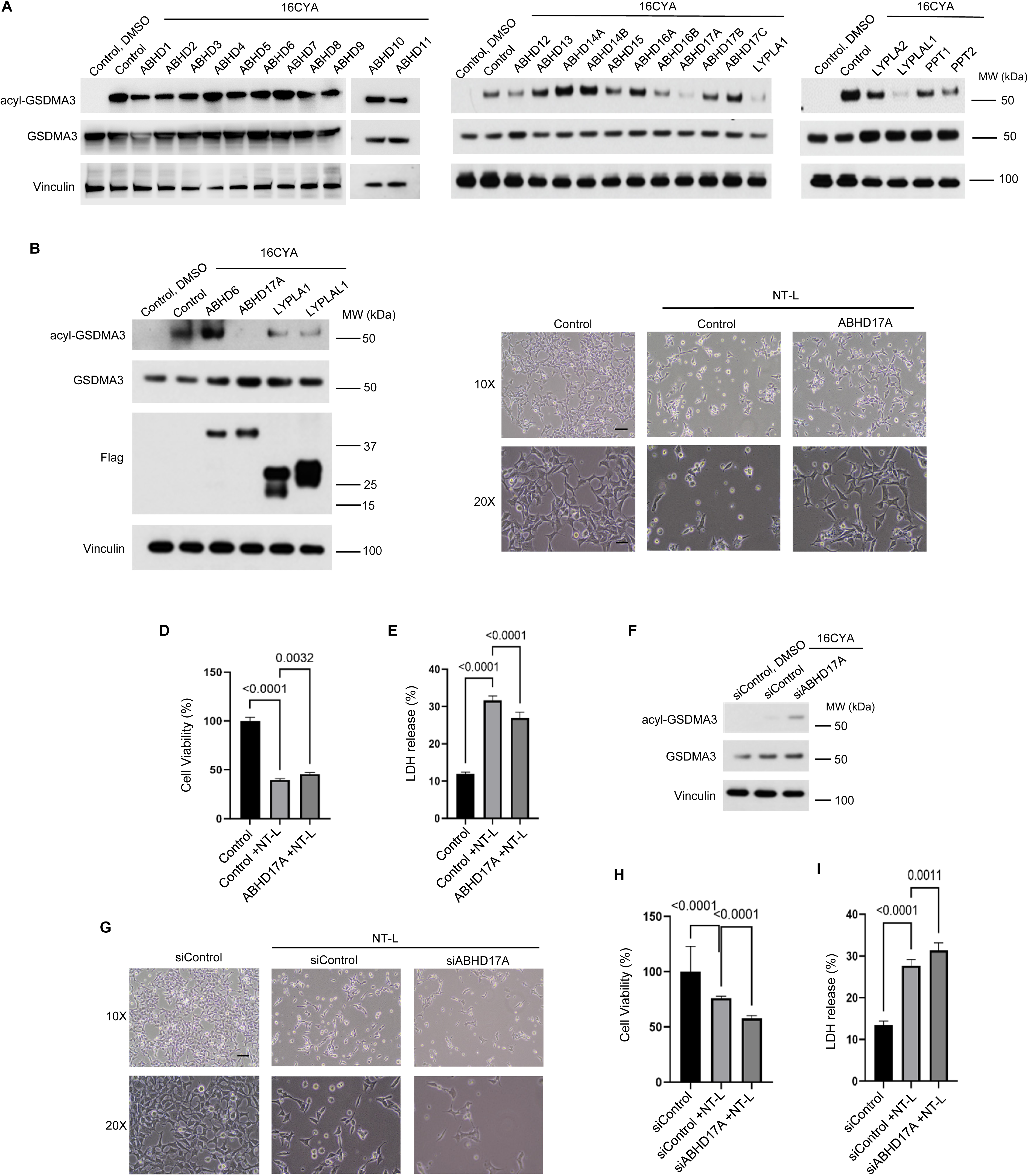
ABHD17A regulates GSDMA3 deacylation and pyroptosis. (A) Screen of 26 deacylating enzymes including ABHDs and APTs on GSDMA3 acylation in transfected HEK293 cells; (B) GSDMA3 acylation levels in HEK293 cells expressing ABHD6, ABHD17A, LYPLA1 or LYPLAL1; (C) The representative images of HEK293 cells expressing GSDMA3 NT-L with or without ABHD17A co-transfection; (D and E) Cell viability (D) and LDH release (E) in HEK293 cells expressing GSDMA3 NT-L with or without ABHD17A co-transfection; (F) ABHD17A siRNA knockdown promotes GSDMA3 acylation; (G) The representative images of HEK293 cells expressing GSDMA3 NT-L with control siRNA (siControl) or siRNA against ABHD17A (siABHD17A); (H and I) Cell viability (H) and LDH release (I) in HEK293 cells expressing GSDMA3 NT-L with control siRNA (siControl) or siRNA against ABHD17A (siABHD17A).

## Discussion

This study characterized the lipid modification of GSDMA using a combination of approaches, including click chemistry, in vitro acylation, structural modeling, and cell-based pyroptosis assays. Prior studies on palmitoylation regulation of gasdermin have focused on GSDMD and its N-terminal C191/C192 (human/mouse) residue as the major site responsible for GSDMD palmitoylation^14–17, 23^. However, C191 is in the “finger” region of the N-terminal domain and is not conserved among different gasdermin proteins. Our analysis not only provided strong evidence that GSDMA also undergoes fatty acylation but identified several specific cysteine residues involved in the “palm” region and demonstrated the functional consequences of their modification. Among the N-terminal cysteine residues, C37 is highly conserved among gasdermin proteins. Furthermore, our structural modeling revealed that palmitates attached to these cysteines are embedded in the membrane and, interestingly, are adjacent to the conserved hydrophobic anchor region identified in gasdermin N-terminal domains, which may provide a molecular mechanism underlying N-terminal lipid acylation of gasdermin family members that facilitates member anchoring and pore formation that is distinct from C191 palmitoylation of GSDMD.

Our study revealed that the recombinant GSDMA and GSDMD proteins can directly attach palmitic acid to themselves without relying on palmityl acyltransferase enzymes. This finding of mammalian gasdermin proteins undergoing fatty acylation *in vitro* by direct interaction with palmitoyl-CoA may open up new avenues of research and potential therapeutic targeting opportunities. Further investigation could focus on identifying the specific amino acid residues within the N-terminal domain of gasdermin that facilitate palmitate transfer. Detailed structural studies and biochemical assays examining how these residues form a catalytic pocket and interact with palmitoyl-CoA could also provide additional evidence of whether the gasdermin proteins may possess “enzyme-like” catalytic activity capable of auto-acylation. Recent reports have shown that palmitoylation of Cys191 at the N-terminus of GSDMD is mediated by palmityl acyltransferases, including ZDHHC5, 7, and 9. Although Cys191 is not conserved among gasdermin proteins, it remains possible that palmityl acyltransferases can also be involved in the acylation of GSDMA. Additionally, multiple potential mechanisms, such as auto-acylation or trans-acylation mediated by ZDHHC enzymes, may fine-tune the regulation of gasdermin lipid modification during pyroptosis.

Our work underscores the potential of targeting GSDMA *S*-acylation for therapeutic applications. We demonstrated that the fatty acylation of GSDMA is dynamically regulated within cells and identified several enzymes that may de-acylate GSDMA, including ABHD17A. This raises the interesting question: Is the acylation-deacylation cycle of GSDMA regulated during skin inflammation, such as infections caused by including group A Streptococcus (GAS)? ^7–10, 32^ Further studies of their functions in vivo could potentially lead to the development of novel therapeutic strategies.

## Methods

### Chemicals

Tris(2-carboxyethyl)phosphine hydrochloride (TCEP) (Cat. #C4706, Sigma Aldrich), Copper(II) sulfate (Cat. #496130, Sigma-Aldrich), Tris[(1-benzyl-1H-1,2,3-triazol-4- yl)methyl]amine (TBTA) (Cat. # 678937, Sigma-Aldrich), chemical probes (Click Chemistry Tools), Streptavidin Agarose beads (Cat. # SA10004, Life Technologies), cOmplete EDTA-free protease inhibitors cocktail (Cat. #05892791001, Roche), Phosphatase inhibitor cocktail (Cat. #P0044, Sigma-Aldrich), Alkyne Palmitoyl-Coenzyme A (Cat. #15968, Cayman), GSDMA recombinant protein (Cat. # TP319050, Origene), GSDMD recombinant protein (Cat. #CSB-EP009956HU, COSMO BIO USA).

### Cell culture and transfection

Human Embryonic Kidney 293A cell line (HEK293A) and human HaCat keratinocyte cell line (HaCaT) were procured from ATCC and cultivated in high-glucose Dulbecco’s modified Eagles media (DMEM) supplied by Life Technologies. 10% (v/v) fetal bovine serum FBS (Thermo/Hyclone, Waltham, MA) and 1x Penicillin/streptomycin (P/S) (GE Health Life Sciences HyClone Laboratories, Logan, UT) were added as supplements to DMEM. None of the cell lines cultured in the study is listed in the database of commonly misidentified cell lines maintained by ICLAC. All cell lines are free of mycoplasma contamination. Expression constructs were transfected into cells using jetPRIME transfection reagent (Polyplus transfection) or Lipofectamine 3000 (Thermo Fisher).

### shRNA knockdown and quantitative RT-PCR

pLKO based lentiviral constructs shRNAs against ZDHHC5 (TRCN0000160468 and TRCN0000166569), ZDHHC7 (TRCN0000275634 and TRCN0000275633), and ZDHHC9 (TRCN0000137411 and TRCN0000137668, purchased from Sigma, St. Louis, MO, USA) were transfected along with the packaging plasmids into growing HEK293T cells. Viral supernatants were collected 48 hours after transfection, and target cells were infected in the presence of polybrene and underwent selection with puromycin for 4-5 days. Total RNA was extracted using the RNeasy mini kit (Qiagen, Cat#74104) or Trizol according to the manufacturer’s instructions. cDNA was synthesized from 2 μg of total RNA using a high-capacity cDNA reverse transcription kit (Life Technologies Cat# 4368814) with random primers, according to the manufacturer’s protocol. Gene expression was quantified using PowerUp SYB Green Master Mix kit (Life Technologies A25777) in the Applied Biosystems system and normalized to β-actin. The following primers were used: ZDHHC5 (Forward: AACTGTATTGGTCGCCGGAAC, and Reverse:

AACACACCCATAATGTGGGCT), ZDHHC7 (Forward: CTGACCGGGTCTGGTTCATC, and Reverse: CATGACGAAAGTCACCACGAA), ZDHHC9 (Forward: CCCAGGCAGGAACACCTTTT, and Reverse: CCGAGGAATCACTCCAGGG), and β-actin (Forward: CACCATTGGCAATGAGCGGTTC, and Reverse: AGGTCTTTGCGGATGTCCACGT).

### Cell labeling and click reaction

HEK293A cells were transfected with either empty vectors or various plasmid constructs. Twenty-four hours after transfection, the cell culture media was discarded. The cells were then washed twice with PBS to remove any remaining fetal bovine serum (FBS) from the DMEM medium. Next, the cells were treated with DMEM supplemented with 10% dialyzed FBS (DFBS) and 1x penicillin/streptomycin (P/S) for two hours. Following this treatment, the cells underwent labeling with chemical probes or DMSO for a duration of six hours. After the labeling, the cells were washed twice with cold PBS and then lysed using a lysis solution composed of 50 mM TEA-HCl (pH 7.4), 150 mM NaCl, 1% Triton X-100, 0.1% SDS, 1 mM PMSF, and a 1x EDTA-free complete protease inhibitor cocktail (Roche). Protein concentration was determined using the BCA method.

A click reaction was performed for one hour at room temperature in 100 µL of lysis buffer containing: 100 µg protein, 100 µM biotin-azide, 1 mM tris(2-carboxyethyl)phosphine hydrochloride (TCEP), 100 µM Tris[(1-benzyl-1H-1,2,3-triazol-4-yl)methyl]amine (TBTA), and 1 mM CuSO4. Following overnight precipitation at -80 °C with nine volumes of cold methanol, the proteins were recovered by centrifugation at 17,000 x g for ten minutes at 4 °C. The precipitates were first suspended in 75 µL of suspension buffer (PBS, 0.05% Tween-20, and 2% SDS). This suspension was then diluted with 350 µL of washing buffer (PBS with 0.05% Tween-20). An aliquot of 75 µL of the diluted suspension was reserved as input, denatured with 6X SDS-PAGE sample buffer, and boiled at 95 °C for 10 minutes. The denatured samples were designated as the input. The remaining 350 µL of the labeled protein suspension was incubated with 30 µL of streptavidin agarose suspension (Life Technologies) for two hours at room temperature with rotation. After two hours, unattached proteins were carefully removed by centrifugation at 500 x g for five minutes at 4 °C following rotation. The protein-bound streptavidin agarose beads were washed three times with PBST (PBS with 0.05% Tween-20). The bound proteins were then eluted for ten minutes at 95 °C using an elution buffer containing 10 mM EDTA (pH 8.2) and 95% formamide. Finally, proteins were resolved by SDS-PAGE after denaturing the samples with 6X SDS-PAGE sample buffer and boiling at 95 °C for 10 minutes post-elution, which were considered as acylated proteins.

### *In vitro* acylation assay

Recombinant His-GSDMA or His-GSDMD protein (500ng) was incubated with the indicated concentrations of alkyne palmitoyl-CoA (Cayman Chemical) for 30 minutes or for the indicated time during the kinetic assay in 50 mM MES buffer (pH 6.4). Our initial experiments using 1 µM of alkyne palmitoyl-CoA showed that both GSDMA and GSDMD proteins reacted with alkyne palmitoyl-CoA. After testing the dost-dependent reactions using 0, 0.1, 1, 2.5, 5, 10 µM of alkyne palmitoyl-CoA, 2.5 µM was selected as the concentration for conducting the time-course reaction, as this concentration produced the highest reaction velocity. For the time-course reaction, the recombinant GSDMA protein was incubated with 2.5 µM of alkyne palmitoyl-CoA for 0, 1, 3, 10, 30, and 50 minutes, and a reaction time of 10 minutes was selected for the kinetic assay, and all the assays were repeated at least three times. For the temperature-dependent assay, 500 nM GSDMA was incubated with 3 µM palmitoyl-CoA at 25, 30, 33, 37, and 42 °C for 30

minutes. For GSDMA concentration-dependent reaction, 100, 200, 300, 400, and 500 nM GSDMA were incubated with 3 µM palmitoyl-CoA at 37 °C for 30 minutes. After the reaction of GSDMA/GSDMD recombinant proteins with alkyne palmitoyl-CoA, CuSO4/TBTA/TCEP/Biotin-Azide master mix was added to the reaction buffer, achieving the following final concentrations: CuSO4 100 mM, TBTA 10 mM, TCEP 100 mM and Biotin-Azide 10 mM. Samples were then incubated for 1 hour at room temperature, followed by SDS-PAGE electrophoresis. Biotinylated GSDMA or GSDMD protein was detected by streptavidin-HRP, and total protein was detected by anti-His antibody (ThermoFisher Scientific, MA1-135). Band intensities obtained from streptavidin blots were quantified using ImageJ (NIH).

### Western Blotting

The cells were lysed using RIPA buffer that had been supplemented with Roche phosphatase and protease inhibitors. Following denaturation at 95 °C for five minutes, lysates were deposited onto 4-12% Bis-Tris polyacrylamide gel. For the SDS-PAGE, NuPAGE MES running reagent (Invitrogen) was utilized. Subsequently, the proteins were transferred to Millipore polyvinylidene fluoride (PVDF) membranes. Following blocking with 5% non-fat milk, the membranes were subjected to incubation with primary and secondary antibodies conjugated to HRP. To detect the protein expression signal, The PVDF membrane was incubated with enhanced chemiluminescent (ECL) horseradish peroxidase (HRP) substrate (ThermoFisher Scientific) and exposed to autoradiographic film (LBSCIENTIFIC INC). Antibody information: anti-GSDMA (CST, #49307S, 1:1000), anti-FLAG M2 (Cat. #F1804, Sigma Aldrich, 1:5000), anti-HA (CST, #3724S, 1:1000), anti-Vinculin (Santa Cruz, sc-73614, 1:3000).

### Immunofluorescent staining

Cells were seeded on glass cover slips in 6-well plate. After culturing for 48 hours, cells were washed with cold PBS for 3 times and fixed with 4% paraformaldehyde for 5 minutes, blocked for 1 hour, and incubated overnight at 4 °C with primary antibody diluted in blocking buffer. Slides were then incubated in Alexa Fluor-conjugated secondary antibodies (Invitrogen) at indicated dilutions in blocking buffer for 2 hours at room temperature before being mounted using a mounting medium with DAPI (EMS). Images were captured by Leica TCS-NT 4D confocal microscope.

### Cell death and viability assays

HaCat cells were seeded into 96-well plates and transfected with GSDMA3-NT-L and GSDMA3-NT-L-3CA plasmids using Lipofectamine 3000. Control cells received a Lipofectamine transfection mixture without plasmid. After 24 hours, cells were stained with propidium iodide (PI) and Hoechst 33342 (Life Technologies, Carlsbad, CA) to assess viability. A staining solution was prepared in 1× PBS, containing 2.5 μM PI and 20 μM Hoechst 33342. 20 μL of this solution was added per well, and plates were incubated at 37 °C with 5% CO₂ for 60 minutes. Cell death was quantified using a Celigo image cytometer (Nexcelom Bioscience). The total and dead cell counts were determined, and the percentage of PI+ cells was calculated and plotted using GraphPad Prism (v.10.0.0). Fluorescent images were captured using the Celigo image cytometer to visualize cell death in HaCat cells. Propidium iodide (PI) staining (red) highlights dead cells, while Hoechst 33342 staining (blue) marks cell nuclei. Cell viability in transfected HEK293 cells was assessed by PrestoBlue™ HS Cell Viability Reagents (ThermoFisher, A13262) according to the manufacturer’s instructions. Absorbance at 570 nm and 600 nm was measured using a BioTek Synergy Neo2 microplate reader, and the data was processed by the difference between 570 nm and 600 nm.

### LDH release measurements

Lactate dehydrogenase (LDH) is a stable cytosolic enzyme found in various cell types. When the plasma membrane is disrupted, LDH is rapidly released into the culture medium, making it a useful marker for cytotoxicity. LDH release was measured using the LDH-Glo Cytotoxicity Assay kit (Promega, J2380), following the manufacturer’s instructions. Cell culture media from different treatments were harvested. A 5 µl sample of the medium was diluted in 45 µl of storage buffer. The LDH detection reagent was then added at a 1:1 ratio. Wells without cells served as negative controls to establish the background of the culture medium. A series of LDH standards (0, 0.5, 1, 2, 4, 8, 16, and 32 mU/ml) were used as positive controls. The reaction was incubated for 30 minutes at room temperature, and luminescence was recorded using a PerkinElmer EnVision plate reader.

### Structural modeling of palmitoylated gasdermin

The structure of palmitoylated GSDMA3 in a lipid bilayer was generated based on a previously reported cryoEM structure of the GSDMA3 pore protein (PDB code 6CB8)^29^. The Modeller program^33^ was used to add coordinates for four residues in each subunit of the 27-mer pore that were not resolved in the cryoEM structure (residues 66-69). CHARMM-GUI membrane builder^34, 35^ was used to model palmitoyl groups on residues C37, C202, and C225 in each subunit, as well as the membrane lipid bilayer context, with no lipids located in the pore center. The lipid bilayer was generated with phosphatidylcholine (POPC) lipids, corresponding to the primary lipid component used for gasdermin experimental characterization in Ruan et al.^29^, and membrane positioning was based on the GSDMA3 pore structure (6CB8) in the Orientations of Membrane Proteins (OPM) database^36^. The system was solvated with TIP3P water^37^ and neutralized with a concentration of 150 mM NaCl. Structural minimization was performed with Amber18^38^, using the CHARMM36m force field^39^. Multiple stages of minimization were performed, with 5000 steepest descent steps each: (i) Initial minimization of water and counter ions with a harmonic constraint of 25 kcal/mol*Å^2^, (ii) Subsequent minimization of water and counter ions with a decreased harmonic constraint of 5 kcal/mol*Å^2^, (iii) Two rounds of minimization of all components of the system with a 5 kcal/mol*Å^2^ harmonic restraint applied exclusively to the protein and attached palmitoyl groups, (iv) Minimization of all components of the system without restraints. Structural figures were generated using PyMOL (Schrodinger, LLC).

### Statistical methods

The sample size was not predetermined statistically. The experiments were not randomized. We carried out biochemical studies at least three independent times. At least three independent experiments that we presented representative images for were completed successfully. The analysis didn’t exclude any samples. When conducting experiments and analyzing the results, the researchers weren’t blinded to the allocation. All results are expressed as means ± s.d. and analyzed by variance analysis (ANOVA) to determine *p* values; *p* < 0.05 was considered statistically significant.

## Supporting Information

Additional experimental details and materials, including GSDMA3 acylation using recombinant proteins and cell culture-based assays.

## Acknowledgment

This work was supported by grants from the National Institutes of Health, R01DK127180 and R01DK127207 to J.M. and R01CA238270 and R01DK107651 to X.W. and in part by a Melanoma Research Alliance (MRA) dermatology postdoctoral fellowship to Z.T. We also thank the members of Wu lab and Mao Lab for helpful discussions and comments on the manuscript. We are grateful to Riya Samanta (University of Maryland), Helder Ribeiro-Filho (Brazilian Biosciences National Laboratory), and S. Saif Hasan (University of Maryland School of Medicine) for input regarding the molecular dynamics simulations and structural modeling.

## Author Contributions

ZT, XW, and JM conceived and designed the study. ZT, RPT, JZ and XH performed the experiments. ZT, RPT, XW and JM analyzed and interpreted the data. MC and BGP performed structural modeling. ZT and JM wrote the manuscript. XW and JM supervised the study.

---

**Figure S1.**
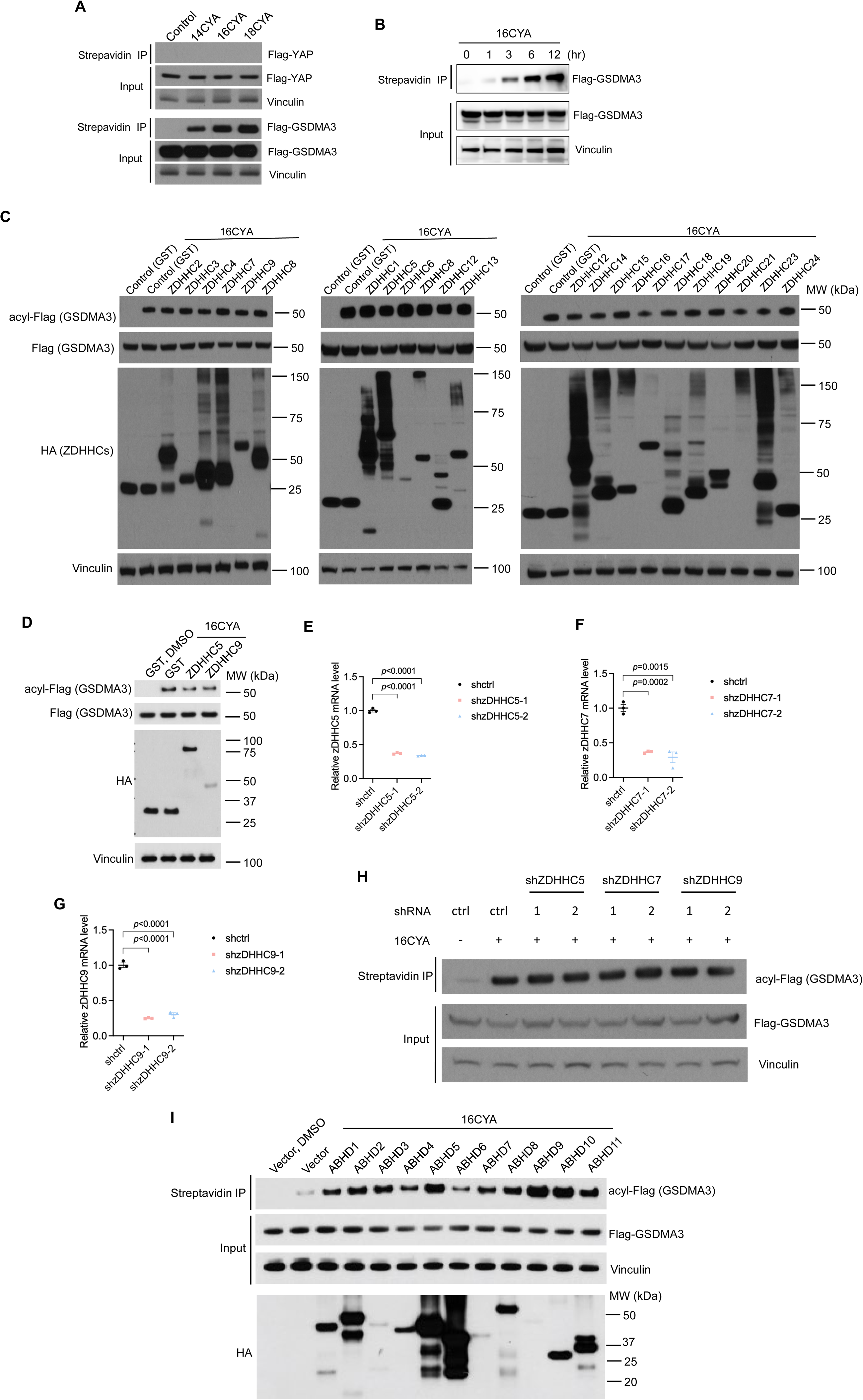
Regulation of GSDMA3 acylation in transfected HEK293 cells. (a) GSDMA3 but not YAP undergoes S-acylation in transfected HEK293 cells measured by different chain length fatty acid probes. Strepavidin immunoprecipitation (IP) detects the acylated proteins. (b) GSDMA3 acylation in HEK293 cells labeled with 16-carbon palmitoylation (16CYA) probe for different durations (0, 1, 3, 6, and 12 hours). (c) GSDMA3 acylation levels in HEK293 cells expressing various ZDHHC enzymes, including ZDHHC5, ZDHHC7, and ZDHHC9 (highlighted in red). (d) GSDMA3 acylation levels in HEK293 cells expressing ZDHHC5 and ZDHHC9. Expression of ZDHHC5, ZDHHC9 and control GST proteins was detected by antidoy against HA tag. (e-g) Relative mRNA levels of ZDHHC5, ZDHHC7, and ZDHHC9 following shRNA knockdown in HEK293 cells (h) GSDMA3 acylation levels in HEK293 cells expressing shRNAs against ZDHHC5, ZDHHC7, and ZDHHC9. (i) GSDMA3 acylation levels in HEK293 cells expressing various ABHD enzymes.

**Figure S2.**
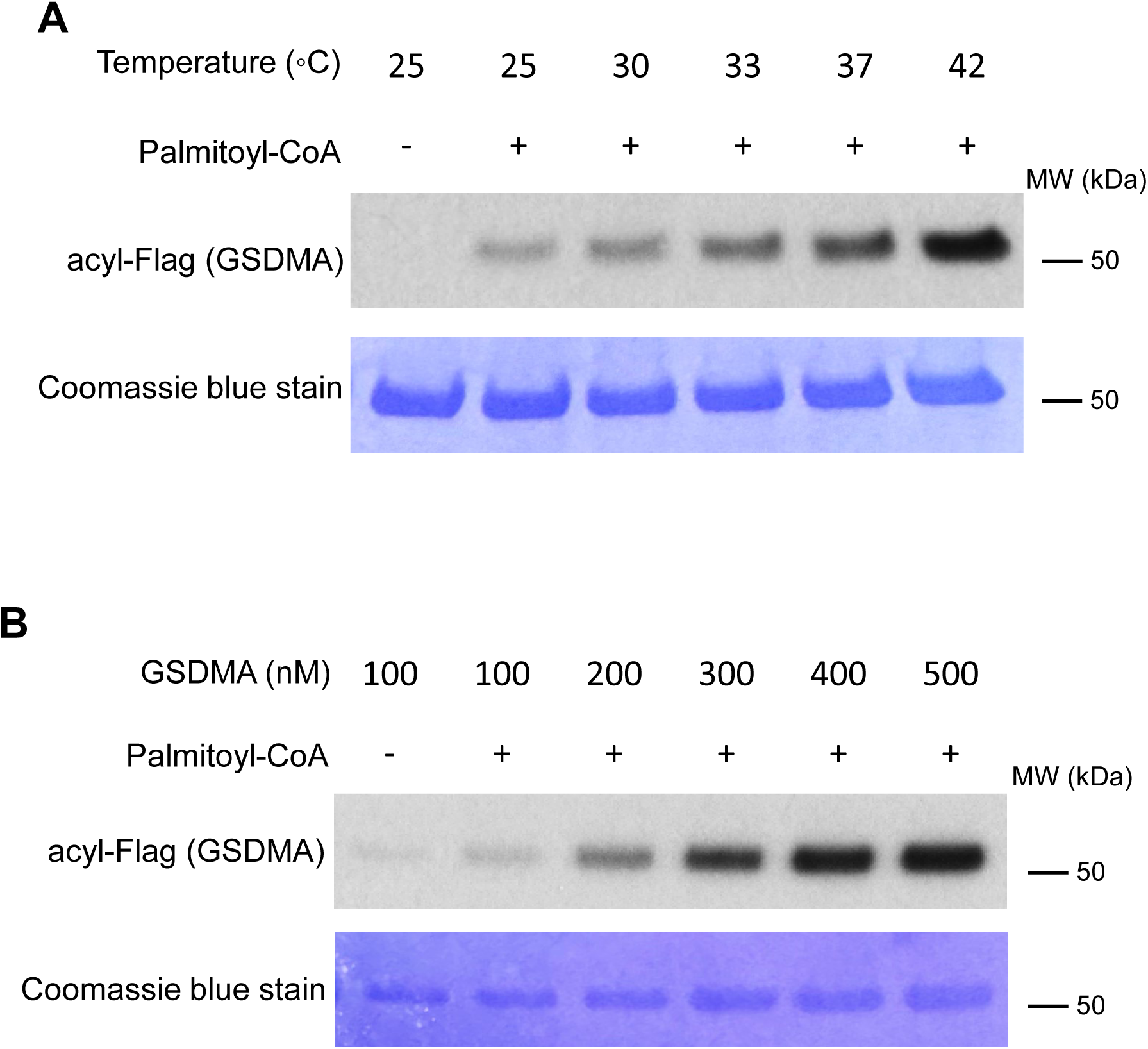
In vitro *S*-acylation of GSDMA. (a) Acylation of GSDMA with 3 µM palmitoyl-CoA at 25, 30, 33, 37, and 42 °C for 30 minutes. (b) Acylation of 100, 200, 300, 400, and 500 nM GSDMA with 3 µM palmitoyl-CoA at 37 °C for 30 minutes.

## References

1. Shi J, Gao W, Shao F. Pyroptosis: Gasdermin-Mediated Programmed Necrotic Cell Death. Trends Biochem Sci. 2017;42(4):245–54. Epub 20161205. doi: 10.1016/j.tibs.2016.10.004. PubMed PMID: 27932073.

2. Broz P, Pelegrin P, Shao F. The gasdermins, a protein family executing cell death and inflammation. Nat Rev Immunol. 2020;20(3):143–57. Epub 20191105. doi: 10.1038/s41577-019-0228-2. PubMed PMID: 31690840.

3. Xia S, Hollingsworth LRt, Wu H. Mechanism and Regulation of Gasdermin-Mediated Cell Death. Cold Spring Harb Perspect Biol. 2020;12(3). Epub 20200302. doi: 10.1101/cshperspect.a036400. PubMed PMID: 31451512; PMCID: PMC7050592.

4. Liu X, Xia S, Zhang Z, Wu H, Lieberman J. Channelling inflammation: gasdermins in physiology and disease. Nat Rev Drug Discov. 2021;20(5):384–405. Epub 20210310. doi: 10.1038/s41573-021-00154-z. PubMed PMID: 33692549; PMCID: PMC7944254.

5. Liu X, Lieberman J. Inflammasome-independent pyroptosis. Curr Opin Immunol. 2024;88:102432. Epub 20240613. doi: 10.1016/j.coi.2024.102432. PubMed PMID: 38875738.

6. Chen KW, Broz P. Gasdermins as evolutionarily conserved executors of inflammation and cell death. Nat Cell Biol. 2024;26(9):1394–406. Epub 20240826. doi: 10.1038/s41556-024-01474-z. PubMed PMID: 39187689.

7. LaRock DL, Johnson AF, Wilde S, Sands JS, Monteiro MP, LaRock CN. Group A Streptococcus induces GSDMA-dependent pyroptosis in keratinocytes. Nature. 2022;605(7910):527-31. Epub 20220511. doi: 10.1038/s41586-022-04717-x. PubMed PMID: 35545676; PMCID: PMC9186297.

8. Deng W, Bai Y, Deng F, Pan Y, Mei S, Zheng Z, Min R, Wu Z, Li W, Miao R, Zhang Z, Kupper TS, Lieberman J, Liu X. Streptococcal pyrogenic exotoxin B cleaves GSDMA and triggers pyroptosis. Nature. 2022;602(7897):496-502. Epub 20220202. doi: 10.1038/s41586-021-04384-4. PubMed PMID: 35110732; PMCID: PMC9703647.

9. Zhao A, Kirkby M, Man SM. Streptococcus makes the cut: Gasdermin A-induced pyroptosis. Cell Host Microbe. 2022;30(4):410–2. doi: 10.1016/j.chom.2022.03.003. PubMed PMID: 35421330.

10. Symmank J, Jacobs C, Schulze-Spate U. Suicide signaling by GSDMA: a single-molecule mechanism for recognition and defense against SpeB-expressing GAS. Signal Transduct Target Ther. 2022;7(1):153. Epub 20220507. doi: 10.1038/s41392-022-01011-0. PubMed PMID: 35525909; PMCID: PMC9079061.

11. Martin BR, Wang C, Adibekian A, Tully SE, Cravatt BF. Global profiling of dynamic protein palmitoylation. Nature methods. 2012;9(1):84–9.

12. Gottlieb CD, Linder ME. Structure and function of DHHC protein S-acyltransferases. Biochemical Society Transactions. 2017;45(4):923–8.

13. Hu L, Tao Z, Wu X. Insights into auto-S-fatty acylation: targets, druggability, and inhibitors. RSC Chem Biol. 2021;2(6):1567–79. Epub 20210825. doi: 10.1039/d1cb00115a. PubMed PMID: 34977571; PMCID: PMC8637764.

14. Du G, Healy LB, David L, Walker C, El-Baba TJ, Lutomski CA, Goh B, Gu B, Pi X, Devant P, Fontana P, Dong Y, Ma X, Miao R, Balasubramanian A, Puthenveetil R, Banerjee A, Luo HR, Kagan JC, Oh SF, Robinson CV, Lieberman J, Wu H. ROS-dependent S-palmitoylation activates cleaved and intact gasdermin D. Nature. 2024;630(8016):437-46. Epub 20240410. doi: 10.1038/s41586-024-07373-5. PubMed PMID: 38599239; PMCID: PMC11283288.

15. Balasubramanian A, Hsu AY, Ghimire L, Tahir M, Devant P, Fontana P, Du G, Liu X, Fabin D, Kambara H, Xie X, Liu F, Hasegawa T, Xu R, Yu H, Chen M, Kolakowski S, Trauger S, Larsen MR, Wei W, Wu H, Kagan JC, Lieberman J, Luo HR. The palmitoylation of gasdermin D directs its membrane translocation and pore formation during pyroptosis. Sci Immunol. 2024;9(94):eadn1452. Epub 20240412. doi: 10.1126/sciimmunol.adn1452. PubMed PMID: 38530158; PMCID: PMC11367861.

16. Liu Z, Li S, Wang C, Vidmar KJ, Bracey S, Li L, Willard B, Miyagi M, Lan T, Dickinson BC, Osme A, Pizarro TT, Xiao TS. Palmitoylation at a conserved cysteine residue facilitates gasdermin D-mediated pyroptosis and cytokine release. Proc Natl Acad Sci U S A. 2024;121(29):e2400883121. Epub 20240709. doi: 10.1073/pnas.2400883121. PubMed PMID: 38980908; PMCID: PMC11260154.

17. Zhang N, Zhang J, Yang Y, Shan H, Hou S, Fang H, Ma M, Chen Z, Tan L, Xu D. A palmitoylation-depalmitoylation relay spatiotemporally controls GSDMD activation in pyroptosis. Nat Cell Biol. 2024;26(5):757–69. Epub 20240327. doi: 10.1038/s41556-024-01397-9. PubMed PMID: 38538834.

18. Sun Z, Hornung V. A critical role for palmitoylation in pyroptosis. Mol Cell. 2024;84(12):2218–20. doi: 10.1016/j.molcel.2024.05.023. PubMed PMID: 38906113; PMCID: PMC7616718.

19. Zhang N, Yang Y, Xu D. Emerging roles of palmitoylation in pyroptosis. Trends Cell Biol. 2024. Epub 20241108. doi: 10.1016/j.tcb.2024.10.005. PubMed PMID: 39521664.

20. Stine L, Humphries F. Gasdermin D palmitoylation: to cleave or not to cleave? Trends Immunol. 2024;45(6):403–5. Epub 20240516. doi: 10.1016/j.it.2024.05.001. PubMed PMID: 38760304.

21. Hu JJ, Liu X, Xia S, Zhang Z, Zhang Y, Zhao J, Ruan J, Luo X, Lou X, Bai Y, Wang J, Hollingsworth LR, Magupalli VG, Zhao L, Luo HR, Kim J, Lieberman J, Wu H. FDA-approved disulfiram inhibits pyroptosis by blocking gasdermin D pore formation. Nat Immunol. 2020;21(7):736–45. Epub 20200504. doi: 10.1038/s41590-020-0669-6. PubMed PMID: 32367036; PMCID: PMC7316630.

22. Humphries F, Shmuel-Galia L, Ketelut-Carneiro N, Li S, Wang B, Nemmara VV, Wilson R, Jiang Z, Khalighinejad F, Muneeruddin K, Shaffer SA, Dutta R, Ionete C, Pesiridis S, Yang S, Thompson PR, Fitzgerald KA. Succination inactivates gasdermin D and blocks pyroptosis. Science. 2020;369(6511):1633-7. Epub 20200820. doi: 10.1126/science.abb9818. PubMed PMID: 32820063; PMCID: PMC8744141.

23. Jiang X, Zhang X, Cai X, Li N, Zheng H, Tang M, Zhu J, Su K, Zhang R, Ye N, Peng J, Zhao M, Wu W, Yang J, Ye H. NU6300 covalently reacts with cysteine-191 of gasdermin D to block its cleavage and palmitoylation. Sci Adv. 2024;10(6):eadi9284. Epub 20240207. doi: 10.1126/sciadv.adi9284. PubMed PMID: 38324683; PMCID: PMC10849585.

24. Zhuang Z, Gu J, Li BO, Yang L. Inhibition of gasdermin D palmitoylation by disulfiram is crucial for the treatment of myocardial infarction. Transl Res. 2024;264:66–75. Epub 20230927. doi: 10.1016/j.trsl.2023.09.007. PubMed PMID: 37769810.

25. Ding J, Wang K, Liu W, She Y, Sun Q, Shi J, Sun H, Wang DC, Shao F. Pore-forming activity and structural autoinhibition of the gasdermin family. Nature. 2016;535(7610):111-6. Epub 20160608. doi: 10.1038/nature18590. PubMed PMID: 27281216.

26. Johnson AG, Wein T, Mayer ML, Duncan-Lowey B, Yirmiya E, Oppenheimer-Shaanan Y, Amitai G, Sorek R, Kranzusch PJ. Bacterial gasdermins reveal an ancient mechanism of cell death. Science. 2022;375(6577):221-5. Epub 20220113. doi: 10.1126/science.abj8432. PubMed PMID: 35025633; PMCID: PMC9134750.

27. Jennings BC, Linder ME. DHHC protein S-acyltransferases use similar ping-pong kinetic mechanisms but display different acyl-CoA specificities. J Biol Chem. 2012;287(10):7236–45. Epub 20120113. doi: 10.1074/jbc.M111.337246. PubMed PMID: 22247542; PMCID: PMC3293542.

28. Chen B, Niu J, Kreuzer J, Zheng B, Jarugumilli GK, Haas W, Wu X. Auto-fatty acylation of transcription factor RFX3 regulates ciliogenesis. Proc Natl Acad Sci U S A. 2018;115(36):E8403-E12. Epub 20180820. doi: 10.1073/pnas.1800949115. PubMed PMID: 30127002; PMCID: PMC6130365.

29. Ruan J, Xia S, Liu X, Lieberman J, Wu H. Cryo-EM structure of the gasdermin A3 membrane pore. Nature. 2018;557(7703):62-7. Epub 20180425. doi: 10.1038/s41586-018-0058-6. PubMed PMID: 29695864; PMCID: PMC6007975.

30. Lin DT, Conibear E. ABHD17 proteins are novel protein depalmitoylases that regulate N-Ras palmitate turnover and subcellular localization. Elife. 2015;4:e11306. Epub 20151223. doi: 10.7554/eLife.11306. PubMed PMID: 26701913; PMCID: PMC4755737.

31. 31. Won SJ, Cheung See Kit M, Martin BR. Protein depalmitoylases. Crit Rev Biochem Mol Biol. 2018;53(1):83–98. Epub 20171214. doi: 10.1080/10409238.2017.1409191. PubMed PMID: 29239216; PMCID: PMC6009847.

32. Brouwer S, Rivera-Hernandez T, Curren BF, Harbison-Price N, De Oliveira DM, Jespersen MG, Davies MR, Walker MJ. Pathogenesis, epidemiology and control of Group A Streptococcus infection. Nature Reviews Microbiology. 2023;21(7):431–47.

33. Webb B, Sali A. Protein structure modeling with MODELLER. Methods in molecular biology. 2014;1137:1–15. doi: 10.1007/978-1-4939-0366-5_1. PubMed PMID: 24573470.

34. Jo S, Kim T, Im W. Automated builder and database of protein/membrane complexes for molecular dynamics simulations. PLoS One. 2007;2(9):e880. Epub 20070912. doi: 10.1371/journal.pone.0000880. PubMed PMID: 17849009; PMCID: PMC1963319.

35. Jo S, Kim T, Iyer VG, Im W. CHARMM-GUI: a web-based graphical user interface for CHARMM. J Comput Chem. 2008;29(11):1859–65. doi: 10.1002/jcc.20945. PubMed PMID: 18351591.

36. Lomize MA, Lomize AL, Pogozheva ID, Mosberg HI. OPM: orientations of proteins in membranes database. Bioinformatics. 2006;22(5):623–5. Epub 20060105. doi: 10.1093/bioinformatics/btk023. PubMed PMID: 16397007.

37. Jorgensen WL, Chandrasekhar J, Madura JD, Impey RW, Klein ML. Comparison of simple potential functions for simulating liquid water. The Journal of chemical physics. 1983;79(2):926–35.

38. Case D, Ben-Shalom I, Brozell S, Cerutti D, Cheatham III T, Cruzeiro V, Darden T, Duke R, Ghoreishi D, Gilson M, Gohlke H, Goetz A, Greene D, Harris R, Homeyer N, Huang Y, Izadi S, Kovalenko A, Kurtzman T, Lee T, LeGrand S, Li P, Lin C, Liu J, Luchko T, Luo R, Mermelstein D, Merz K, Miao Y, Monard G, Nguyen C, Nguyen H, Omelyan I, Onufriev A, Pan F, Qi R, Roe D, Roitberg A, Sagui C, Schott-Verdugo S, Shen J, Simmerling C, Smith J, Salomon-Ferrer R, Swails J, Walker R, Wang J, Wei H, Wolf R, Wu X, Xiao L, York D, Kollman P. AMBER 2018. University of California, San Francisco. 2018.

39. Huang J, Rauscher S, Nawrocki G, Ran T, Feig M, de Groot BL, Grubmuller H, MacKerell AD, Jr. CHARMM36m: an improved force field for folded and intrinsically disordered proteins. Nat Methods. 2017;14(1):71–3. Epub 20161107. doi: 10.1038/nmeth.4067. PubMed PMID: 27819658; PMCID: PMC5199616.

